# Nuclear-cytoplasmic balance: whole genome duplications induce elevated organellar genome copy number

**DOI:** 10.1101/2021.06.08.447629

**Authors:** Matheus Fernandes Gyorfy, Emma R. Miller, Justin L. Conover, Corrinne E. Grover, Jonathan F. Wendel, Daniel B. Sloan, Joel Sharbrough

## Abstract

The plant genome is partitioned across three distinct subcellular compartments: the nucleus, mitochondria, and plastids. Successful coordination of gene expression among these organellar genomes and the nuclear genome is critical for plant function and fitness. Whole genome duplication events (WGDs) in the nucleus have played a major role in the diversification of land plants and are expected to perturb the relative copy number (stoichiometry) of nuclear, mitochondrial, and plastid genomes. Thus, elucidating the mechanisms whereby plant cells respond to the cytonuclear stoichiometric imbalance that follow WGDs represents an important yet underexplored question in understanding the evolutionary consequences of genome doubling. We used droplet digital PCR (ddPCR) to investigate the relationship between nuclear and organellar genome copy numbers in allopolyploids and their diploid progenitors in both wheat and *Arabidopsis*. Polyploids exhibit elevated organellar genome copy numbers per cell, largely preserving the cytonuclear stoichiometry observed in diploids despite the change in nuclear genome copy number. To investigate the timescale over which cytonuclear stoichiometry may respond to WGD, we also estimated organellar genome copy number in *Arabidopsis* synthetic autopolyploids and in a haploid-induced diploid line. We observed corresponding changes in organellar genome copy number in these laboratory-generated lines, indicating that at least some of the cellular response to cytonuclear stoichiometric imbalance is immediate following WGD. We conclude that increases in organellar genome copy numbers represent a common response to polyploidization, suggesting that maintenance of cytonuclear stoichiometry is an important component in establishing polyploid lineages.

**Significance Statement:** Whole genome duplications (WGD) have the potential to alter the stoichiometric balance between nuclear and organellar genomes. We used two separate diploid-polyploid complexes to show that plant cells with WGD exhibit elevated mitochondrial and plastid genome copy numbers, both immediately in lab-generated lines and in natural polyploids.

## Introduction

Plant cells require coordinated communication among nuclear, mitochondrial, and plastid genomes to perform necessary metabolic functions (Greiner and Bock, 2013; Wang *et al*., 2020). Many of the key enzyme complexes in mitochondria and plastids (including chloroplasts) comprise a mix of nuclear-encoded and cytoplasmically encoded subunits that must be assembled in specific ratios (Forsythe *et al*., 2019). Whole genome duplication events (WGDs) in the nucleus may potentially disrupt this cytonuclear communication by altering the relative copy number (stoichiometry) of these genomes (Sharbrough *et al*., 2017).

Whole genome duplication events, whether by allopolyploidization or autopolyploidization, have been pervasive throughout the evolutionary history of angiosperms (Wood *et al*., 2009; Jiao *et al*., 2011; Wendel, 2015; Ruprecht *et al*., 2017; One Thousand Plant Transcriptomes Initiative, 2019). Studies in polyploid plants have uncovered dramatic impacts of ploidy on nuclear genome evolution and plant biology (Wendel *et al*., 2018; Doyle and Coate, 2019; Bomblies, 2020; Fox *et al*., 2020; Van de Peer *et al*., 2021). For example, WGDs can lead to genome shock where dormant transposable elements regain their movement throughout the genome. Other important impacts include short-term global transcriptomic and epigenetic changes affecting expression, and longer-term nuclear genome diploidization.

The process of diploidization, in which duplicate gene copies are lost following WGD, appears to be non-random (De Smet *et al*., 2013), with selection acting to preserve “gene balance” between sets of functionally interacting loci (Coate *et al*., 2011; Song *et al*., 2020). The importance of nuclear gene balance has been further supported by observations that associate partial genomic redundancy (e.g., aneuploidy) with increased deleterious effect relative to WGD (Birchler and Veitia, 2012; Shi *et al*., 2020). Polyploidy is also expected to affect stoichiometry and scaling relationships among other cellular components because of the broadly observed positive correlation between nuclear genome size and cell size (Beaulieu *et al*., 2008; Doyle and Coate, 2019).

Although fewer studies have investigated the effects of WGDs on cytonuclear stoichiometry, various lines of evidence indicate that perturbations from nuclear genome doubling may be mitigated by correlated changes in the cytoplasm. For example, numerous polyploid angiosperms exhibit an increase in chloroplasts per cell compared to related diploids, although the extent of this increase and whether it scales with the change in nuclear genome copy number varies substantially across taxa (Bingham, 1968; Krishnaswami and Andal, 1978; Beversdorf, 1979; Ellis and Leech, 1985; Butterfass, 1987; Jacobs and Yoder, 1989; Ho *et al*., 1990; Singsit and Ozias-Akins, 1992; Warner and Edwards, 1993; Beck *et al*., 2003; Ewald *et al*., 2009; Murti *et al*., 2012; He *et al*., 2021). While presence of systematic increases in mitochondrial number in plant polyploid species is less clear, there is evidence that mitochondrial count, size, and function can scale with cell size in eukaryotes (Preuten *et al*., 2010; Rafelski *et al*., 2012; Cole, 2016; Miettinen and Björklund, 2017). Patterns regarding balance between the organellar and nuclear genomes are further complicated by variation in the number of genome copies per organelle among cell types, which has been shown to scale negatively organelle number per cell in some cases (Cattolico, 1979; Dean and Leech, 1982; Lamppa *et al*., 1992; Preuten *et al*., 2010; Shen *et al*., 2019). Attempts to quantify the copy number ratio for nuclear vs. organellar genomes in polyploids suggest that increased amounts of plastid and mitochondrial DNA may stabilize cytonuclear gene balance following WGDs, although these studies have not always detected the same levels of compensation (Whiteway and Lee, 1977; Bowman, 1986; Dean and Leech, 1982; Oberprieler *et al*., 2019; Coate *et al*., 2020).

A recent study of *Arabidopsis thaliana* autopolyploids simultaneously investigated changes in both plastid-nuclear and mitochondrial-nuclear genomic stoichiometry (Coate *et al*., 2020). Specifically, the authors used quantitative PCR (qPCR) to show a reduced ratio of plastid genome copies per nuclear genome copy in polyploids relative to diploids, whereas the mitochondrial-nuclear ratio was maintained. Coate and colleagues (Coate *et al*., 2020) also found that polyploids maintained diploid-like expression ratios between organellar genomes and the nuclear genes that encode mitochondrial- and plastid-targeted proteins, but the mitochondria and plastids appeared to do so by different routes. That is, nuclear-encoded, plastid targeted genes and plastid-encoded genes were both proportionally downregulated in polyploids relative to the rest of the nuclear genome. By contrast, nuclear-encoded, mitochondrially targeted genes and mitochondrial-encoded genes both maintained stable expression or were even upregulated in polyploids relative to the rest of the nuclear genome (Coate *et al*., 2020). Together, these studies provide mounting evidence that at least some cytonuclear stoichiometric compensation occurs after an increase in nuclear ploidy, but we have only limited information on how this compensation may differ between plastids and mitochondria. In addition, it is unclear why levels of compensation may differ across taxa. One possibility is that this relationship depends on the timing of polyploidization, because more ancient polyploids may exhibit an evolved response, whereas new polyploids may only exhibit an immediate developmental response. Thus, researching the differing effects of WGDs on cytonuclear stoichiometry is critical to understanding the origins and evolution of polyploid organisms.

To explore changes in cytonuclear stoichiometry accompanying WGD, we selected two well-studied diploid-polyploid complexes, wheat and *Arabidopsis*. A repeated history of polyploidization and hybridization in both complexes make them well-suited models for understanding the consequences of genome doubling for cytonuclear stoichiometry. *Triticum turgidum* is an allotetraploid (2n = 4x) that resulted from a genome merger between two diploid wheat lineages (represented by extant relatives *T. urartu* (2n = 2x) and *Aegilops speltoides* (2n = 2x)) roughly 0.5 – 1.0 MY ago (Marcussen *et al*., 2014; Avni *et al*., 2017; Maccaferri *et al*., 2019). A second allopolyploidization event, in which pollen from diploid *Aegilops tauschii* (2n = 2x) fertilized *T. turgidum*, gave rise to allohexaploid *T. aestivum* (2n = 6x) *ca*. 10,000 years ago (Huang *et al*., 2002; Marcussen *et al*., 2014; Li *et al*., 2015; Sandve *et al*., 2015; El Baidouri *et al*., 2017). A similar, but evolutionarily independent history of WGD resulted in allotetraploid *Arabidopsis suecica* (2n = 4x) and is thought to have originated from an allopolyploidization event between an unreduced *A. thaliana* (2n = 2x) ovule and tetraploid *A. arenosa* (4n = 4x) pollen *ca*. 16,000 years ago (Jakobsson *et al*., 2006; Novikova *et al*., 2017; Burns *et al*., 2021). Together, these independent allopolyploids provide a useful comparative context to investigate the evolved cytonuclear stoichiometric responses to WGD. To assess the immediate effects of WGDs, we took advantage of the availability of synthetic (i.e., lab-generated) lines of *A. thaliana* that differ in ploidy. These included diploid *A. thaliana* Columbia-0 (Col-0) lines that underwent colchicine treatment to produce autotetraploid and auto-octoploid lines (Robinson *et al*., 2018), as well as the natural tetraploid *A. thaliana* Warschau (Wa-1) accession (4n = 4x) that underwent haploid induction to reduce ploidy and generate a synthetic diploid (Ravi and Chan, 2010).

We used droplet digital PCR (ddPCR) to accurately and precisely measure nuclear and organellar genome copy numbers in diploid vs. polyploid leaf tissue from plants raised in a common garden. In contrast to the relative quantification offered by qPCR, ddPCR provides an estimate of the absolute copy numbers by partitioning samples into thousands of nanoreactor droplets that are individually analyzed for the presence or absence of amplification (hence, digital), and an absolute copy number can be inferred using a Poisson distribution (Hindson *et al*., 2011). Overall, we observed clear signs of cytonuclear stoichiometry compensation based on increased organellar genome copy numbers in plants with higher nuclear ploidy.

## Results

### Variation in organellar genome copy number during development in Aegilops speltoides

Before assessing how organellar genome copy number is affected by a change in ploidy, we first measured copy number variation across leaves collected at three different developmental sampling times (based on tiller development; see Methods) in a single diploid species (*Aegilops speltoides*). We tested the extent to which development influenced organellar genome copy number, which in turn allowed us to time DNA extractions in a way that would limit developmental effects on cytonuclear stoichiometry when comparing across different species. On average, we estimated that *Aegilops speltoides* leaf tissue harbors approximately 50 to 100 times more plastid genome copies per cell (mean +/- SD = 2,402 +/- 963) than mitochondrial genome copies per cell (mean +/- SD = 44 +/-10) (Figure 1). Notably, the two double-copy genes that we used (mitochondrial *atp6* and plastid *ndhB*) as internal controls of organellar genome copy number exhibited counts between 1.8- to 2.0-times higher than those of the single-copy genes, indicating that our ddPCR-based estimates of genome copy number were precise and consistent across markers.

**Figure 1.**
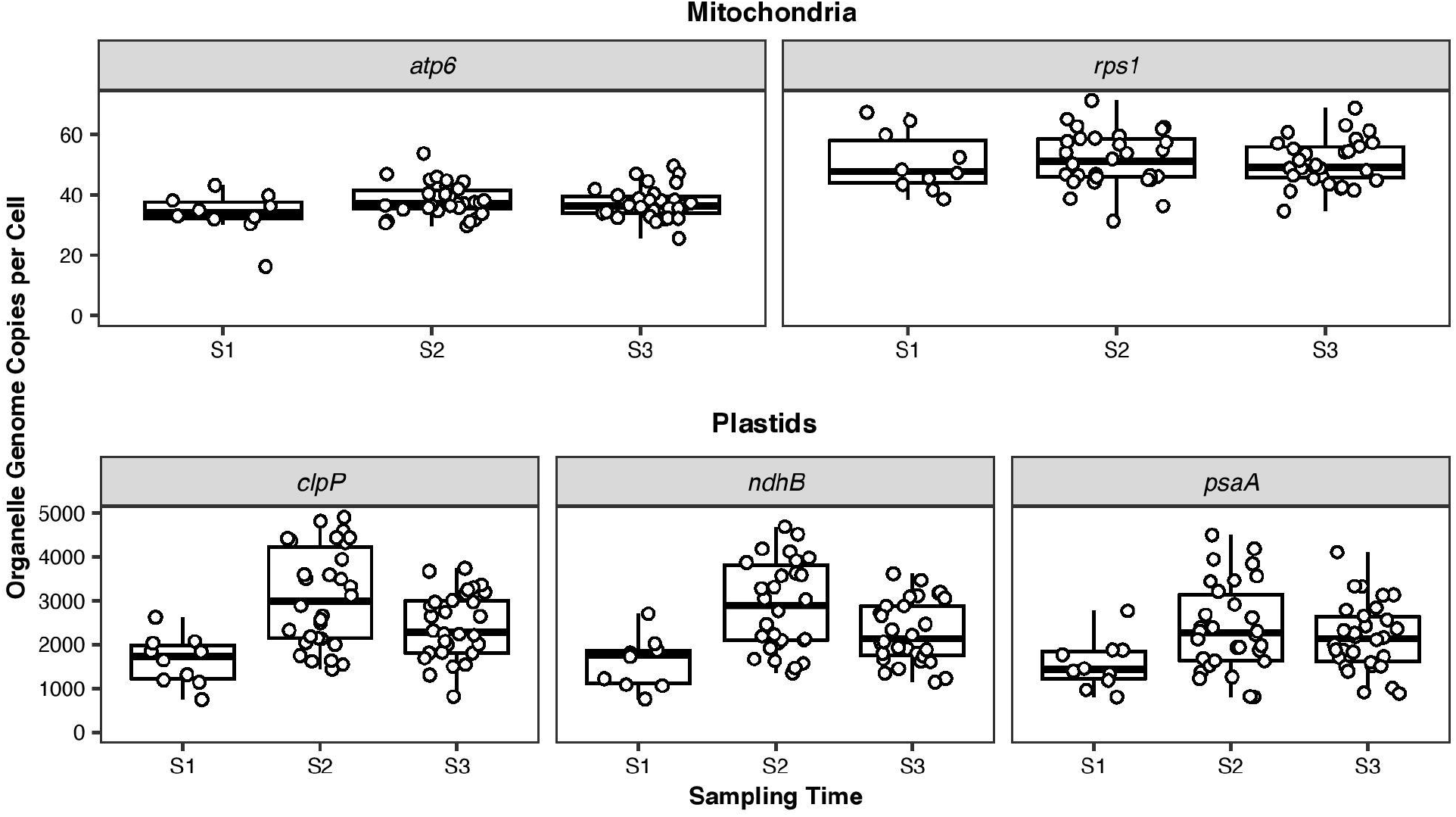
Organellar genome copy numbers remain similar across three developmental sampling times in *Aegilops speltoides*. Organellar genome copy numbers per cell in *Aegilops speltoides* (2x). The y-axis represents the number of organellar copies per cell for each genome marker estimated from ddPCR. The “per cell” calculations are based on nuclear markers (Table S2) and assume that cells are at their standard ploidy and have not yet undergone DNA replication (S phase) or rounds of endoreduplication. Values for *atp6* and *ndhB* were divided by two to account for the fact that they are found in two-copy repeats. The x-axis represents the sampling times for different developmental stages (see Methods).

Organellar genome copy counts remained similar across the three sampling time points. While sampling time was a marginally significant predictor of overall mitochondrial genome copy number (χ^2^ = 7.6637; df = 2; *p* = 0.0217), pairwise comparisons between individual sampling times were not significantly different, reflecting a small effect of developmental stage on mitochondrial copy number. Development had a larger effect on plastid genome copy number (χ^2^ = 38.6319; df = 2; *p* < 0.0001); however, the effect size was still modest in scale as the mean plastid genome copy counts remained within a 2-fold range across sampling times. In light of the developmental variation detected, we standardized a single sampling time for all measurements from both wheat and *Arabidopsis* representatives (see Methods).

### Allopolyploid wheat exhibits elevated organellar genome copy number per cell

We next used allopolyploid wheat species (*T. turgidum* [4x] and *T. aestivum* [6x]) and related diploids (*Aegilops speltoides* and *T. urartu*) grown in a common garden to test whether polyploids exhibit elevated numbers of organellar genomes per cell. We found that species was a significant predictor of genome copy number per cell for both organelles (mitochondrial: χ^2^ = 473.649, df = 3, *p* < 0.0001; plastid: χ^2^ = 186.934, df = 3, *p* < 0.0001), with the polyploids exhibiting 2 to 3 times more organellar genomes than the diploids (Figures 2 and S1). Although the two polyploid species did not differ significantly from each other in mitochondrial counts (*t* = −0.089; df = 33; *p* = 0.93), all pairwise comparisons between diploid and polyploid samples were significantly different (*p* < 0.0001 in all cases). In addition, the two diploids differed significantly from each other (*t* = −5.693; df = 33; *p* < 0.0001), with *T. urartu* (mean +/- SD = 57 +/- 14) exhibiting higher mitochondrial genome counts than *Aegilops speltoides* (mean +/- SD = 37 +/-12). Polyploids also exhibited significantly higher plastid genome copy numbers per cell than diploids (*p* < 0.0001 in all four pairwise comparisons); however, tetraploid *T. turgidum* (mean +/- SD = 4120 +/- 813) and hexaploid *T. aestivum* (mean +/- SD = 4905 +/- 808) did not differ in the number of plastid genomes per cell (Figure 2). The similar counts between the two polyploid species (4x and 6x) may not reflect a lack of compensation in the hexaploid but rather “overcompensation” in the tetraploid species. When we standardized mitochondrial and plastid genome counts and expressed them on a per haploid nuclear genome copy basis (as opposed to per cell based on the known ploidy level in the species), the tetraploid *T. turgidum* had higher values than either of the two diploids, indicating that *T. turgidum* had more than compensated for the doubling of the nuclear genome (Figure S1). In contrast to mitochondrial genome copy number, the diploid wheat species did not differ from one another in terms of plastid genome copy number (*T. urartu* mean +/- SD = 1814 +/- 377; *Aegilops speltoides* mean +/- SD = 1509 +/- 530, *t* = −2.023*, df* = 33*, p =* 0.1025). In sum, these data indicate clear signs of cytoplasmic compensation such that polyploids exhibit elevated organellar genome copy numbers per cell compared to diploids.

**Figure 2.**
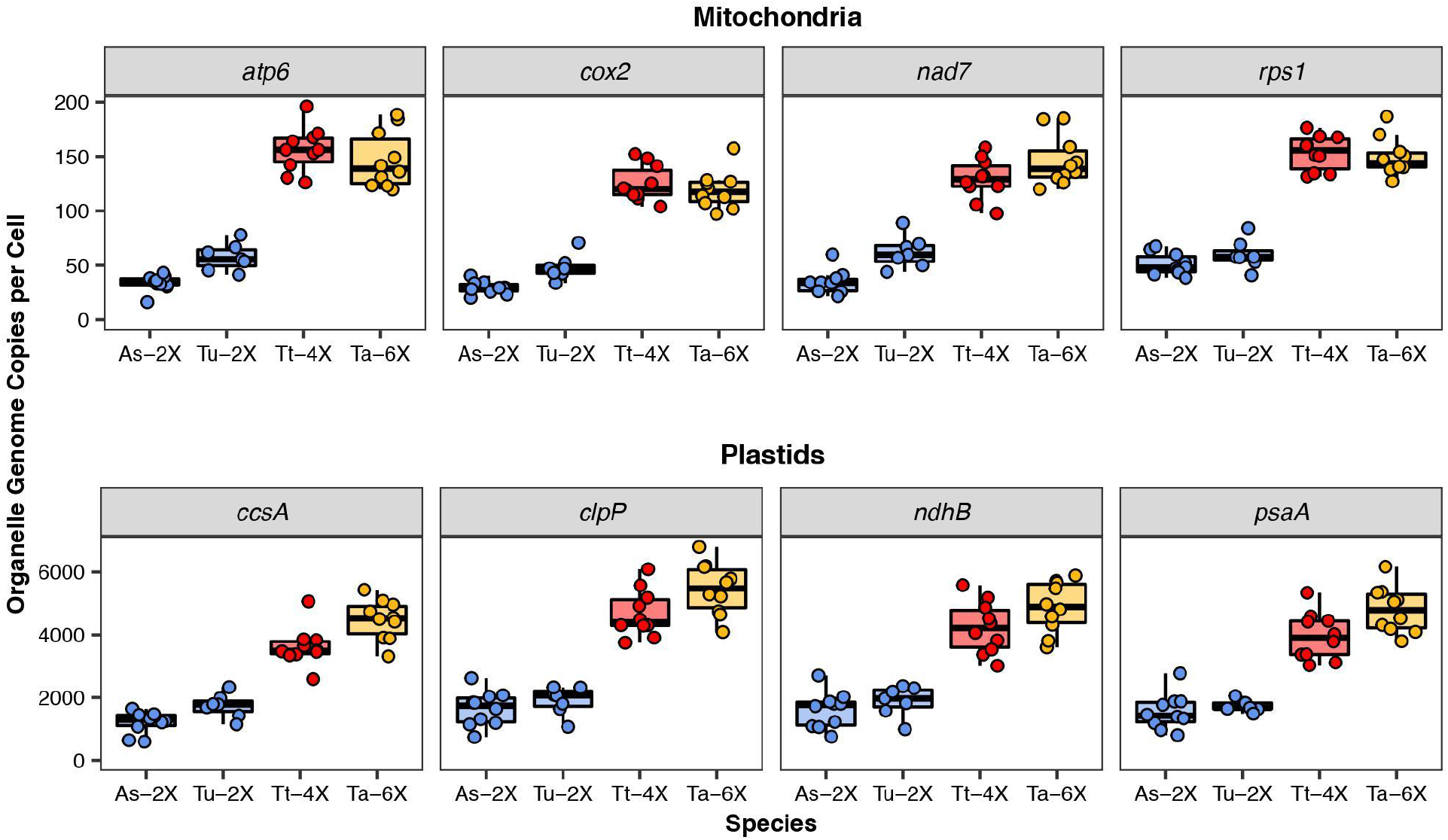
Increased organellar genome copies per cell in wheat allopolyploids. Organellar genome copy numbers per cell in *Aegilops speltoides* (2x, blue), *Triticum urartu* (2x, blue), *Triticum turgidum* (4x, red), and *Triticum aestivum* (6x, yellow). The y-axis represents the number of organellar genome copies per cell for each genome marker estimated from ddPCR. The “per cell” calculations are based on nuclear markers (Table S2) and assume that cells are at their standard ploidy and have not yet undergone DNA replication (S phase) or rounds of endoreduplication. Values for *atp6* and *ndhB* were divided by two to account for the fact that they are found in two-copy repeats.

### Elevated organellar genome copy number in allopolyploid Arabidopsis

To test whether the increase in organellar genome copy number in response to WGD that we observed in polyploid wheat was a general pattern across independent and phylogenetically distant WGDs, we employed a similar analysis for allopolyploid *Arabidopsis suecica* (4x) and its diploid relatives (*A. thaliana* [2x] and *A. arenosa* [2x]). As before, we found that species was a significant predictor of organellar genome copy numbers and that higher nuclear ploidy was associated with an increase in organellar genome copy numbers per cell (mitochondrial: χ^2^ = 25.829, df = 2, *p* < 0.0001; plastid: χ^2^ = 124.1222, df = 2, *p* < 0.0001; Figures 3 and S2). Diploid species exhibited significantly different mitochondrial genome copies per cell from one another (*p* = 0.0008), as *A. thaliana* averaged 10 copies per cell (SD = 2) while *A. arenosa* averaged 15 copies per cell (SD = 6). For the tetraploid, we estimated an average of 26 copies per cell (SD = 8), almost exactly double the average counts between the two diploids. For plastid genome copy counts, we observed 189 copies per cell (SD = 53) in *A. thaliana*, whereas the other diploid, *A. arenosa*, displayed 394 copies per cell (SD = 124). The tetraploid *A. suecica* compensated by more than doubling the average count of both diploid species at 858 copies per cell (SD = 194). Thus, increased organellar genome copy number was directionally similar in the two studied systems.

**Figure 3.**
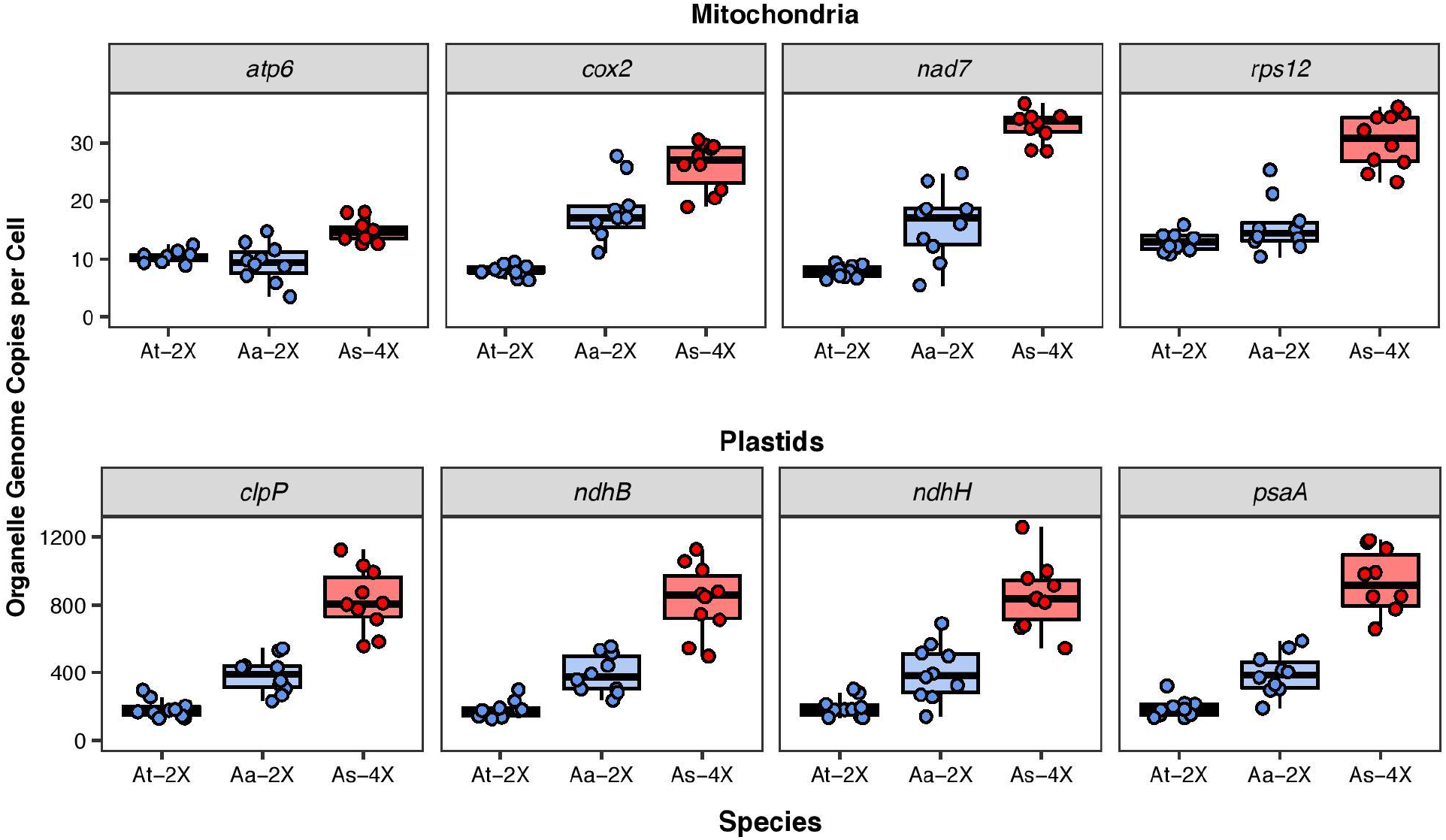
Increased organellar genome copies per cell in an *Arabidopsis* allopolyploid. Organellar genome copy numbers per cell in *Arabidopsis thaliana* (2x, blue), *Arabidopsis arenosa* (2x, blue), and *Arabidopsis suecica* (4x, red). The y-axis represents the number of organellar genome copies per cell for each genome marker estimated from ddPCR. The “per cell” calculations are based on nuclear markers (Table S2) and assume that cells are at their standard ploidy and have not yet undergone DNA replication (S phase) or rounds of endoreduplication. Values for *atp6* and *ndhB* were divided by two to account for the fact that they are found in two-copy repeats.

In addition to species, we found that gene marker was a significant predictor of genome copy number estimates in mitochondria (χ^2^ = 224.722, df = 3, *p* < 0.0001) but not of plastid genome copy numbers (χ^2^ = 0.1205, df = 3, *p* = 0.99). This difference between organelles highlights a general phenomenon observed across multiple comparisons in this study, where there was larger variation among gene markers for mitochondrial genome copy number than for plastid genome copy number. We also observed a significant gene x species interaction in the mitochondrial dataset (χ^2^ = 358.000, df = 6, *p* < 0.0001), indicating that this intragenomic variation differed among species. The patterns observed for *atp6* were particularly interesting. The *A. thaliana* mitochondrial genome has two copies of this gene (Unseld *et al*., 1997), as also indicated by our copy number estimates. By contrast, *A. arenosa* and *A. suecica* exhibit copy number estimates consistent with those obtained from single-copy genes in those species (Figures 3 and S2). Clearly, the observed variation in *atp6* across species relative to other genes contributes to the significant interaction in the mitochondrial dataset, but comparisons among other mitochondrial genes also exhibited differences in relative copy number across species. For example, *cox2* has a higher average copy number than *rps12* in *A. arenosa*, but the opposite is true in *A. thaliana* and *A. suecica* (Figure 3).

### Synthetic autopolyploid Arabidopsis thaliana lines exhibit cytoplasmic compensation as an immediate response to increases in ploidy

Our results from wheat and *Arabidopsis* allopolyploids provided evidence for cytoplasmic compensation in naturally formed polyploid lineages, raising the question of whether this was an evolved response acquired over many generations, or if instead it was an immediate consequence of WGD. To evaluate this, we quantified organellar genome copy number in synthetic autopolyploid lines of *A. thaliana* Col-0 surveyed three generations after synthesis, including both tetraploid (4x) and octoploid (8x) plants (Robinson *et al*., 2018). We observed that ploidy was a highly significant predictor of organellar genome copies per cell in both genomic compartments (mitochondrial: χ^2^ = 188.80, df = 2, *p* < 0.0001; plastid: χ^2^ = 126.7979, df = 2, *p* < 0.0001). With the organellar genome copy numbers roughly doubling with each doubling of nuclear ploidy, these results are consistent with a nearly perfect full compensation model (means of 10, 20, and 40 mitochondrial genome copies per cell in 2x, 4x, and 8x, respectively; means of 189, 404, and 706 plastid genome copies per cell in 2x, 4x, and 8x, respectively; Figure 4). Indeed, when we standardized organellar genome copy counts per haploid nuclear genome copy, there was no effect of ploidy on either mitochondrial or plastid genome copy number estimates (mitochondrial: χ^2^ = 0.9617, df = 2, *p* = 0.62; plastid: χ^2^ = 0.5895, df = 2, *p* = 0.74; Figure S3). The fact that cytoplasmic compensation was detected in lab-generated lines indicates that it can occur immediately with the onset of polyploidy.

**Figure 4.**
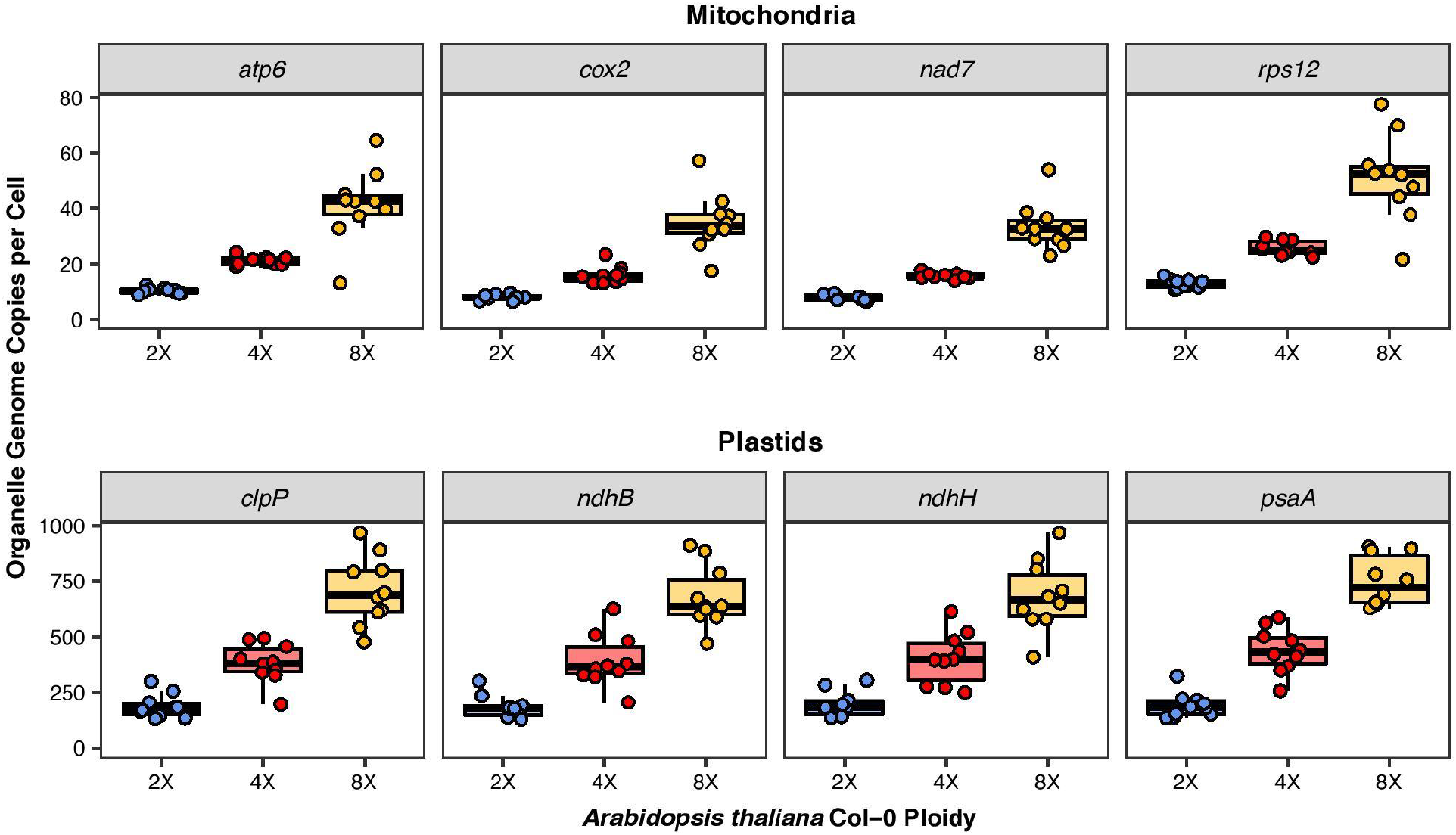
Increased organellar genome copies per cell in an *Arabidopsis* synthetic autopolyploid line. Organellar genome copy numbers per cell in a synthetic polyploid series for *Arabidopsis thaliana* Col-0: 2x (blue), 4x (red), and 8x (yellow). The y-axis represents the number of organellar genome copies per cell for each genome marker estimated from ddPCR. The “per cell” calculations are based on nuclear markers (Table S2) and assume that cells are at their standard ploidy and have not yet undergone DNA replication (S phase) or rounds of endoreduplication. Values for *atp6* and *ndhB* were divided by two to account for the fact that they are found in two-copy repeats.

### Reducing ploidy by haploid-induction in Arabidopsis thaliana Wa-1 results in limited changes in organellar genome copy number

As a corollary to our prediction that polyploids would exhibit elevated organellar genome copies per cell compared to diploids, we also predicted that decreasing ploidy would result in reduced organellar genome copy numbers. To test this prediction, we took advantage of a haploid-induced line of the natural autotetraploid *A. thaliana* Wa-1 accession in which the tetraploid had been converted into a diploid (Ravi and Chan, 2010). Although we observed an overall downward trend in the predicted direction (i.e., diploid Wa-1 exhibited fewer organellar genome copies per cell than tetraploid Wa-1), we did not find a full two-fold reduction in either mitochondrial or plastid genome copy number in response to haploid-induction (Figure 5). The tetraploid:diploid copy number ratio was 1.65 for the mitochondrial genome and 1.14 for the plastid genome. Accordingly, we found that ploidy was a significant predictor of mitochondrial genome copy number (χ^2^ = 10.556, df = 1, *p* = 0.0012), but not of plastid genome copy number (χ^2^ = 0.8718, df = 1, *p* = 0.35). The diploid exhibited significantly more plastid genome copies per haploid nuclear genome copy (*t* = 3.035, df = 18, *p* = 0.0071; Figure S4), allowing us to reject a full two-fold reduction in plastid genome copy number. By contrast, we could not reject a full compensation model for the mitochondrial dataset (χ^2^ = 1.633, df = 18, *p* = 0.12), indicating that a compensatory response is still taking place following this haploid-induction event. Overall, the haploid-induction comparison provides support for cytoplasmic compensation when nuclear ploidy is reduced, but these effects appear weaker than many of the cases of ploidy increases examined.

**Figure 5.**
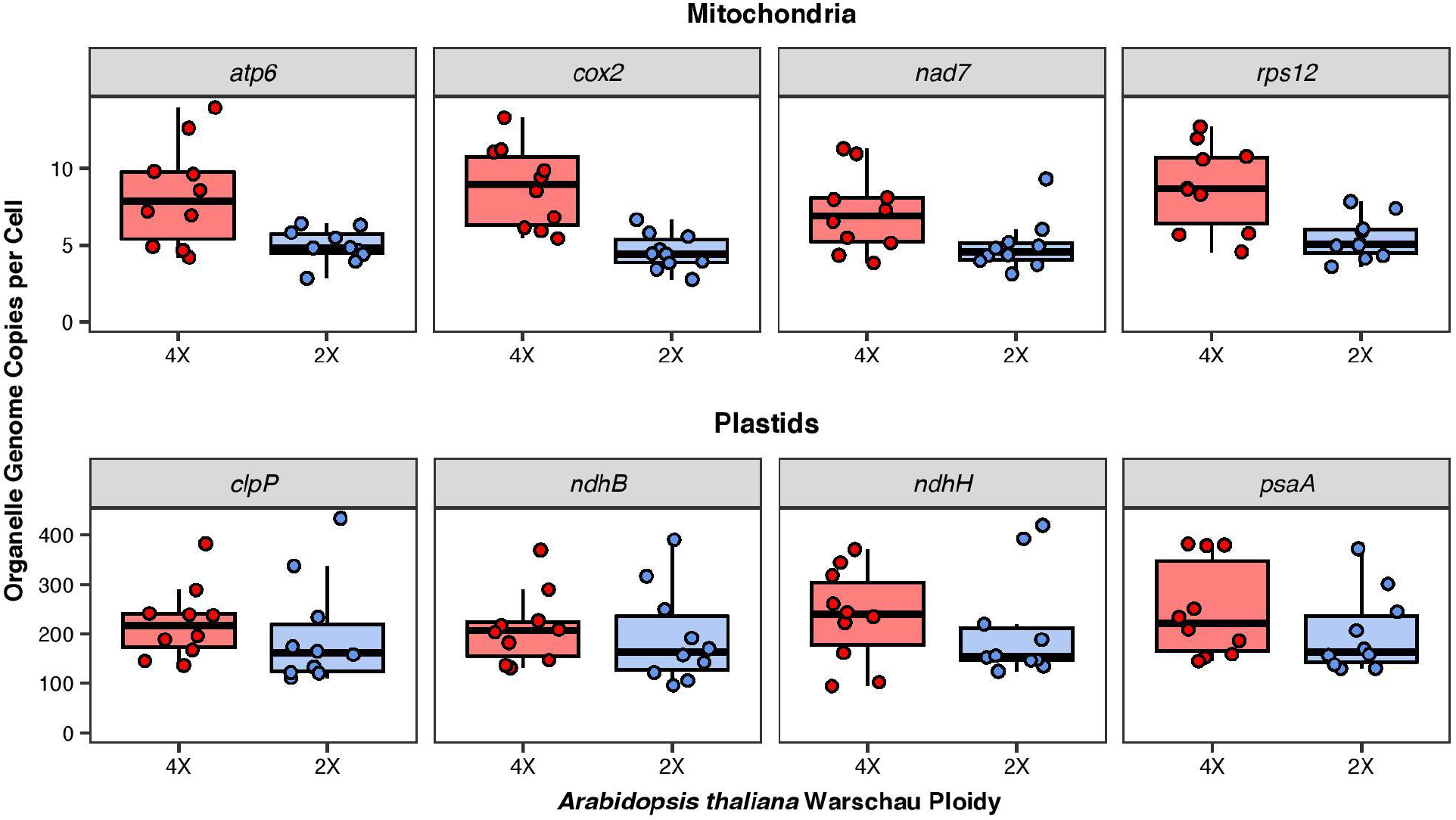
Decreased organellar genome copies per cell in an *Arabidopsis* haploid induction line. Organellar genome copy numbers per cell in *Arabidopsis thaliana* Wa-1 ecotype (4x, red) and a derived haploid induction line (2x, blue). The y-axis represents the number of organellar genome copies per cell for each genome marker estimated from ddPCR. The “per cell” calculations are based on nuclear markers (Table S2) and assume that cells are at their standard ploidy and have not yet undergone DNA replication (S phase) or rounds of endoreduplication. Values for *atp6* and *ndhB* were divided by two to account for the fact that they are found in two-copy repeats.

## Discussion

In polyploids, the organellar genomes exist inside cells whose nuclear genome has suddenly been doubled. The potential for disrupted cytonuclear stoichiometry to negatively affect phenotypes thus exists in nascent polyploid lineages. Here, we tested the extent to which WGDs perturb the balance between nuclear and organellar genomes in two diploid-polyploid species complexes, including both monocot and eudicot taxa. We observed compensation of organellar genome copy number in response to a change in nuclear ploidy in mitochondria and plastids and in both natural and synthetic accessions. We did not find evidence that synthetic accessions exhibited consistently weaker signatures of compensation than natural polyploids, indicating that maintaining cytonuclear stoichiometry represents an important process in the immediate establishment of polyploid lineages.

A number of different mechanisms could explain the organellar genome responses that we observed. First, there is a well established relationship between nuclear genome size and cell size, so polyploids may accommodate more organelles per cell (Marshall *et al*., 2012; Wendel *et al*., 2018; Doyle and Coate, 2019; Bomblies, 2020; Fox *et al*., 2020; Van de Peer *et al*., 2021). Indeed, plant polyploids typically have bigger cells with more plastids per cell (Wendel *et al*., 2018; Doyle and Coate, 2019; Bomblies, 2020; Fox *et al*., 2020; Van de Peer *et al*., 2021). Bigger cell size may trigger an increase in organelle biogenesis (Kawade *et al*., 2013), but our understanding of the complex cellular mechanisms that link genome size, cell size, and organelle number remains incomplete. The relationship between nuclear genome size and number of mitochondria per cell has received less attention in plants, but it is plausible that they exhibit similar scaling relationships as plastids, considering the observed increase in genome copy numbers for both organelles as nuclear ploidy increases (Figures 2–5). The relationship between nuclear genome size and cell size could also serve as an explanation for the large difference in organellar genome copy number between wheat and *Arabidopsis*, in which diploids differ *ca*. 40-fold in genome size (Pellicer and Leitch, 2020). Accordingly, we observed substantially higher copy number estimates in wheat species compared to *Arabidopsis* species of the same ploidy level (Figures 2–5). Whether the correlation between genome size and organellar genome copy number represents a broadly distributed scaling relationship across species remains unclear, but the extensive variation in genome size across angiosperms will allow this hypothesis to be tested.

An additional explanation for the observed cytoplasmic compensation could be a higher number of organelles per unit cell volume in species with an increase in nuclear genome size; however, comparisons across ploidy levels in wheat suggest that polyploids do not increase chloroplast density relative to diploids (Ellis and Leech, 1985; Butterfass, 1987; Warner and Edwards, 1993). It is also possible that the number of genome copies per organelle increases with nuclear ploidy via a denser distribution of genome copies throughout the organelle and/or an increase in the overall size of the organelles. Previous studies have shown that the number of genome copies per chloroplast may differ throughout plant development (Greiner *et al*., 2020) and that chloroplast sizes may both vary and be negatively correlated with cellular chloroplast counts (Kostoff, 1938; Honda *et al*., 1971; Ellis and Leech, 1985), but to our knowledge associations with ploidy have not been evaluated in this context. It seems likely that compensation involves a mixture of these mechanisms whose relative contributions may be revealed through studies coupling microscopy and nucleic acid quantification (Greiner *et al*., 2020) across ploidy levels.

Our calculations of organellar genome copy number “per cell” made the simplifying assumption that all cells were at their respective base ploidy level for the corresponding species or line. Notably, plants can undergo endoreduplication, in which there are one or more rounds of DNA replication without cell division (Joubès and Chevalier, 2000; Edgar *et al*., 2014). It is possible that our data underestimated organellar genome copy numbers per cell if a significant fraction of the leaf cells have experienced endoreduplication. Species with large genomes, including polyploids, have been shown to undergo lower rates of endoreduplication (Barow and Meister, 2003; Pacey *et al*., 2020). Therefore, the difference in the actual average amount of nuclear DNA per cell in diploids vs. polyploids may be lower than implied by their nominal ploidy levels, resulting in a form of immediate compensation in the polyploid species.

Although we saw a general pattern of cytoplasmic compensation, this was not stoichiometrically perfect. For example, tetraploid and hexaploid wheat displayed similar mitochondrial genome copy numbers despite their difference in nuclear genome size. When interpreting such exceptions, it is worth considering that the species we investigated differ from each other in many respects, and some degree of variation among species and lines is expected regardless of ploidy. To this point, the two diploids in our allopolyploid wheat experiment exhibited substantially different organellar genome copy numbers (Figure 2), and the same was true for the two diploids in the allopolyploid *Arabidopsis* experiment (Figure 3). Moreover, when we compared *A. thaliana* Wa-1 and *A. suecica*, both natural tetraploids, the latter had approximately four times as many organellar genome copies (Figures 3 and 5).

Our ddPCR approach allowed us to simultaneously measure organellar genome copy numbers in both plastids and mitochondria in a parallel fashion. Therefore, we could compare and contrast the responses of these two organelle systems. Overall, both organelles responded similarly in terms of compensation following changes in nuclear ploidy, with two possible exceptions. First, hexaploid *T. aestivum* exhibited higher plastid genome copy numbers than tetraploid *T. turgidum*, but not mitochondrial (Figure 2). Second, the synthetic diploid *Arabidopsis thaliana* Wa-1 exhibited significantly fewer genome copies per cell for mitochondria, but not for plastids, compared to its natural tetraploid counterpart, which was in contrast to the allopolyploid wheat results (Figure 5). Given that these two exceptions deviated in opposite directions, there was no general pattern for either organelle system to exhibit a stronger compensatory response. In comparing variation in copy number estimates *within* organellar genomes, we did observe that there was often more heterogeneity among mitochondrial markers than among plastid markers. This phenomenon is consistent with past observations that copy number can vary across the length of the mitochondrial genome in a “fluid” fashion (Preuten et al., 2010; Shen et al., 2019; Wu et al., 2020). The highly recombinational nature of plant mitochondrial genomes is known to make them prone to rapid structural rearrangements, amplification of subgenomic regions, and changes in relative copy numbers (Arrieta-Montiel and Mackenzie, 2011; Gualberto and Newton, 2017).

As noted above, we tested the possibility that lab-generated polyploids would exhibit weaker compensatory responses than more ancient polyploid lineages that had been shaped by a history of natural selection to restore cytonuclear stoichiometry. However, we did not find consistent differences between lab-generated lines and natural polyploids in their degrees of compensation (Figures 2–5). The immediate compensation for cytonuclear stoichiometry that we observed in synthetic *Arabidopsis* lineages coupled with the well documented and widely observed cycle of post-WGD diploidization (Wendel, 2015), sets up the alternative prediction that natural polyploids might exhibit reduced numbers of organellar genome copies per cell compared to synthetic polyploids, as polyploids lose or silence nuclear genes over time. To wit, nuclear genes whose products are targeted to the mitochondria or plastids are among the first to re-diploidize and return to single copy following WGD (De Smet et al. 2013).

In contravention to that expectation, we found that natural polyploids maintained elevated organellar genome copy numbers per cell relative to diploids, with the relatively ancient *T. turgidum* being perhaps the best example, and that the cytonuclear stoichiometry of natural polyploids did not appear to differ from synthetic accessions in any noticeable way. Indeed, *Arabidopsis suecica* exhibited similar or higher organellar genome copy numbers per cell as the synthetic tetraploid (with greater plastid genome copy numbers even than the octoploid), indicating that the immediate cytonuclear stoichiometric compensation does not appear to degrade rapidly post-WGD. It is possible that frequent changes in nuclear ploidy that plants experience in both their cell cycle (endoreduplication) and life cycle (alternation of generations) predispose them to having efficient regulatory responses that maintain cytonuclear stoichiometry following WGDs. The evidently dynamic nature of the compensatory response in synthetic lineages juxtaposed against the persistence of elevated organellar genome copy number across many thousands of generations in natural polyploids highlights the profound and ratchet-like changes that accompany WGD.

Our results demonstrate clear signs of compensation at the DNA level, but further investigation at the RNA and protein levels will also provide insight into cytonuclear function in plant cells after WGD (Oberprieler *et al*., 2019; Coate *et al*., 2020; Yaha *et al*., 2021). Key stoichiometric relationships play out in protein-protein and protein-RNA interactions, creating a need for more transcriptome and proteomic datasets across species of different ploidy levels. Finally, the consistent finding that changes in ploidy resulted in corresponding changes in organellar genome copy number across phylogenetically distinct taxa and multiple independent duplication events means that characterizing the molecular mechanisms responsible for maintaining cytonuclear stoichiometry inside the plant cell represents an important and promising avenue of future investigation.

## Methods

### Seed sources, plant growth conditions, and tissue sampling

Wheat and *Arabidopsis* seeds were obtained from the sources listed in Table S1. Within each diploid-polyploid complex, we grew all plants in parallel and under identical common garden conditions. For the wheat complex, both *T. urartu* and *Aegilops speltoides* were vernalized for 28 days, and *T. aestivum* and *T. turgidum* were vernalized for 7 days at 4°C before being germinated on petri dishes with water-soaked filter paper. Following germination, seedlings were potted in 0.5 gallon pots using Pro-Mix high porosity soil mix (Pro-Mix, Quakertown, USA) with ~5 ml of Osmocote slow-release fertilizer (Scotts Miracle-Gro Company, Marysville, USA) added to the top of each pot. Plants were randomly distributed across growth shelves and grown under a 16-hour day length at 22°C. For *Aegilops speltoides*, we sampled leaves for DNA extraction at three separate developmental times. We sampled (1) the first leaf from a developing tiller whose second leaf just emerged; (2) the second leaf from a tiller whose third leaf had just emerged; and (3) the third leaf from a tiller whose fourth leaf had just emerged. For other species in the wheat complex, we only used samples from the first sampling time point. All *Arabidopsis* species underwent a vernalization period of 10 days at 4°C before being germinated on petri dishes with water-soaked filter paper. We potted seedlings in individual 2.5” square cups using Pro-Mix high porosity soil mix. We randomly distributed plants across growth shelves and grew them under a 16-hour day length at 22°C. For this complex, we waited until a rosette had produced 12 leaves, at which time we sampled the sixth leaf for DNA extraction and ddPCR analysis.

### DNA extractions

We sampled one leaf per individual plant for all species at the sampling times described above, using a razor blade to cut leaves from plants. When leaves were >100 mg, we identified leaf midpoint with respect to length and cut off equal amounts of tissue from both ends until below 100 mg. Leaves were placed in 1.5 mL microcentrifuge tubes and flash frozen in liquid nitrogen. We ground leaves in a liquid nitrogen bath and extracted DNA using the Qiagen DNeasy Plant Mini Kit (Qiagen, Germantown, USA) according to the manufacturer’s instructions. DNA concentrations were quantified using the Qubit dsDNA HS Kit on a Qubit 2.0 (Invitrogen, Carlsbad, USA) and diluted in 10 mM Tris-HCl pH 8.0 to target concentrations for ddPCR (see below).

### Primer development

We designed primers for 12 genes for each diploid-polyploid complex (24 primer pairs in total), four from each genome (i.e., nuclear, mitochondrial, and plastid). Because we used the nuclear primers as a proxy for the number of nuclear genome copies, we ensured that all nuclear primers were single-copy genes in diploids via BLAST of the expected amplicons against the genome assembly. Only amplicons that lacked significant BLAST hits to other genomic regions were utilized in ddPCR estimates of nuclear genome copy number (Table S2). Likewise, we confirmed that allopolyploids had retained all copies of these genes and that primers were a match for all gene copies. For plastid and mitochondrial genomes, we followed a similar approach for three genes each (Table S2), but we also included a fourth gene that was identified as double-copy to serve as an internal control to assess our sensitivity for detecting copy number differences (mitochondrial – *atp6*, plastid – *ndhB)*. We aligned gene sequences from all species using MAFFT v7.470 (Katoh and Standley, 2013), manually designed primers of 18-20 nt in length, and calculated melting temperature and self-compatibility using OligoCalc v3.26 (Kibbe, 2007). The resulting amplicons were between 180 to 210 bp in length which allowed for high efficiency and short replication time during amplification. We confirmed the length of the amplicon for all primers by PCR and gel electrophoresis using the same reaction conditions as were used for ddPCR amplification (see below).

### ddPCR

We performed 20μl ddPCR reactions using 10 μl of BioRad QX200 ddPCR EvaGreen Supermix (BioRad, Hercules, USA), 0.1 μM F primer, 0.1 μM R primer, and 0.1 – 20 ng of total cellular DNA, depending on the genomic compartment and target species. The use of different DNA concentrations across genomic compartments was necessary to prevent saturation of droplet count. For the wheat diploid-polyploid complex, we used 20 ng, 1.0 ng, and 0.1 ng of template DNA for nuclear, mitochondrial, and plastid markers, respectively. For the *Arabidopsis* diploid-polyploid complex, we used 1.0 ng, 0.1 ng, and 0.005 ng of template DNA for nuclear, mitochondrial, and plastid markers, respectively. These concentrations were determined based on the proportion of positive droplets recovered during preliminary primer testing.

We used the Bio-Rad ddPCR system (Bio-Rad, Hercules, USA) to estimate nuclear, mitochondrial, and plastid copy number by generating droplets using a QX200 Droplet Generator according to the manufacturer’s instructions and transferred samples to a 96-well plate. We sealed the plate using the BioRad PX1 PCR Plate Sealer and amplified target gene markers on a Bio-Rad C1000 deep-well thermal cycler. PCR amplification proceeded as follows: a denaturation cycle at 95°C for 5 minutes, followed by 40 cycles of denaturation for 30 seconds at 90°C and annealing/extension at 60°C for 1 minute. Lastly, there was a 5-minute resting period at 4°C, followed by a 5-minute incubation at 90°C, and a final hold at 4°C until droplet reading. We analyzed positive droplet counts using a Bio-Rad QX200 Droplet Reader and the Hex1 channel. Copy number was estimated in Bio-Rad software Quantasoft v1.7, which employs a Poisson distribution to infer the marker copy number in the original DNA sample.

Samples that did not meet quality thresholds were re-run if they met any of the following criteria: (1) failed reaction (low/no positive droplets), (2) reactions with 90% or more positive droplets out of the total droplet count, (3) reactions yielding fewer than 9,000 total droplets, or (4) reactions that were large outliers compared to other samples from the same primer/species combination. Results from re-runs were only included in additional analyses (and original runs excluded) when they disagreed substantially with those from the original runs.

Because the droplet generator processes samples in batches of eight, we designed primer combinations (PCs) that divided batches into two DNA samples with four markers each. All PCs possessed at least one marker gene per genome (Table S3). Results from individual ddPCR samples are provided in Table S3. For each PC, nuclear markers were used to normalize mitochondrial and plastid genome copy numbers from that same PC per cell and per haploid nuclear genome copy. The number of cells sampled for a given PC was estimated by dividing the nuclear marker’s absolute copy number per 20μL reaction by the known ploidy of the sample. For PCs with two nuclear genes, the mean copy number per 20μL between the two markers was used to estimate the number of cells sampled. We then estimated the number of mitochondrial and plastid genomes per cell by multiplying the absolute copies per 20μL reaction by the dilution factor and dividing by the inferred number of cells. We estimated the number of mitochondrial and plastid genome copies per haploid nuclear genome copy by multiplying the absolute copies per 20μL reaction by the dilution factor and dividing by the estimated number of cells and by the known ploidy of the sample. As in the nuclear markers, we used the mean between the two markers as the absolute copy number per 20μL reaction for PCs with either two mitochondrial or two plastid markers. Absolute copy numbers per 20μL reaction of double-copy genes were divided by two to directly compare with single-copy genes.

### Statistical Analyses

All statistical analyses and data visualization were carried out in R v 4.0.3 (R Core Team, 2020). For the *A. speltoides* dataset, We implemented a mixed-effects modeling approach to test whether developmental time point (see above), gene (mitochondria: *atp6, rps 1;* plastid: *ccsA, clpP, ndhB, psaA*), and a development × gene interaction were good predictors of mitochondrial or plastid genome copy numbers per cell and per haploid nuclear genome copy (see above for description of how these values were estimated), with separate models fit for each organelle. A term for plant identity was fit as a random intercept to account for repeated measures on individual plants (Arnqvist, 2020). For allopolyploid comparisons, we employed a similar approach to test whether species (wheat: *Aegilops speltoides, T. urartu, T. turgidum, T. aestivum; Arabidopsis: Arabidopsis thaliana* Col-0, *Arabidopsis arenosa*, *Arabidopsis suecica*), gene (mitochondria: *atp6, cox2, nad7, rps 1* [wheat only], *rps12* [*Arabidopsis* only]; plastid: *ccsA* [wheat only]*, clpP, ndhB, ndhH [Arabidopsis only], psaA*), and an interaction of species × gene were good predictors of mitochondrial or plastid genome copy numbers per cell and per haploid nuclear genome copy. Species was used as a proxy for ploidy in the allopolyploid models because (1) ploidy is confounded with species, and (2) multiple diploid species, but only a single species from each polyploid level, were assayed. We employed a similar framework for synthetic line comparisons (i.e., *Arabidopsis thaliana* Col-0 2x, 4x, 8x; *Arabidopsis thaliana* Wa-1 4x and 2x), except we included a term for ploidy in these models, as all synthetic plant lines were from the same species (*Arabidopsis thaliana*). Mixed-effects models were constructed using the lmer function in the lme4 R package (Bates *et al*., 2007), and the Anova function with type III sums-of-squares computation implemented the car package was used to perform model testing (Fox *et al*., 2007). *Post hoc* pairwise *t* tests were performed using the Satterthwaite approximation of marginal means as implemented by the emmeans package (Satterthwaite, 1946). To account for multiple comparisons, we used the Holm procedure for the Bonferroni correction (Holm, 1979). Results were visualized using ggplot2 (Wickham, 2011). R code, input files, and output are available at https://github.com/mgyorfy/Polyploidy_ddPCR. All statistical results are summarized in Table S4.

## Acknowledgements

This work was funded by the National Science Foundation (IOS-1829176), Colorado State University, and the New Mexico Institute of Mining and Technology. We would like to thank John Anderson for help developing and troubleshooting ddPCR assays and Carol Wilusz and the Colorado State University Molecular Quantification Core for training and access to digital droplet PCR instrumentation. Wheat seeds were obtained from the USDA’s National Small Grains Collection, part of the Germplasm Resources Information Network, and housed at Colorado State University. We thank Scott Haley, Patrick Byrne, and Scott Seifert for advice on wheat accession choices and growth conditions. We thank Jeremy Coate for providing synthetic *Arabidopsis* seeds, Andreas Madlung for *Arabidopsis arenosa* and *Arabidopsis suecica* seeds and advice on *Arabidopsis* growth conditions, and Luca Comai, Adrienne Roeder, Maruthachalam Ravi, and Simon WL Chan for generating synthetic *Arabidopsis* lines.

## FIGURES

**Figure S1.**
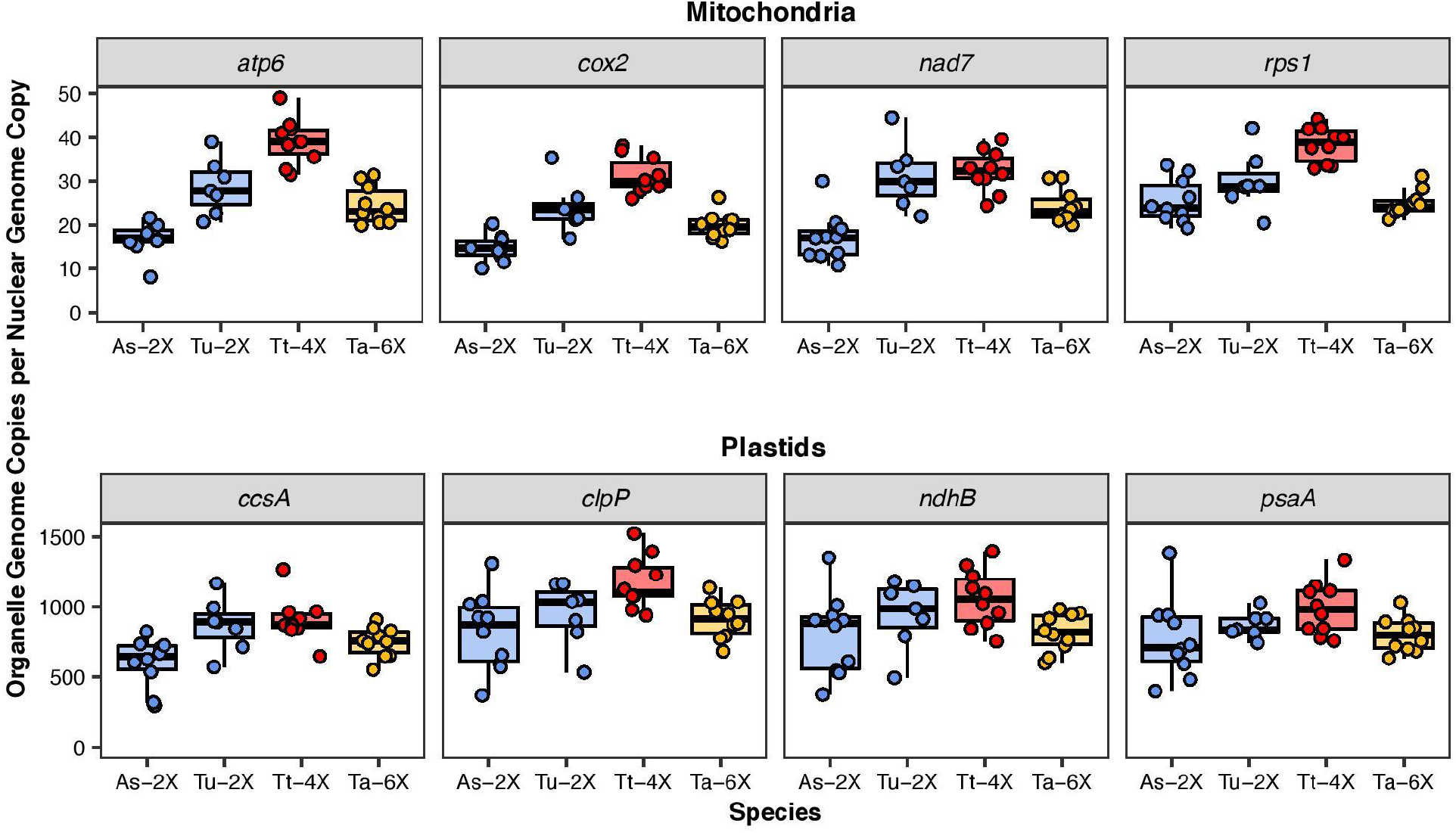
Maintenance of cytonuclear stoichiometric ratios in wheat allopolyploids. Organellar genome copy numbers per haploid nuclear genome copy in *Aegilops speltoides* (2x, blue), *Triticum urartu* (2x, blue), *Triticum turgidum* (4x, red), and *Triticum aestivum* (6x, yellow). The y-axis shows the estimated number of organellar genome copies per haploid nuclear genome copy (see Methods). Values for *atp6* and *ndhB* were divided by two to account for the fact that they are found in two-copy repeats.

**Figure S2.**
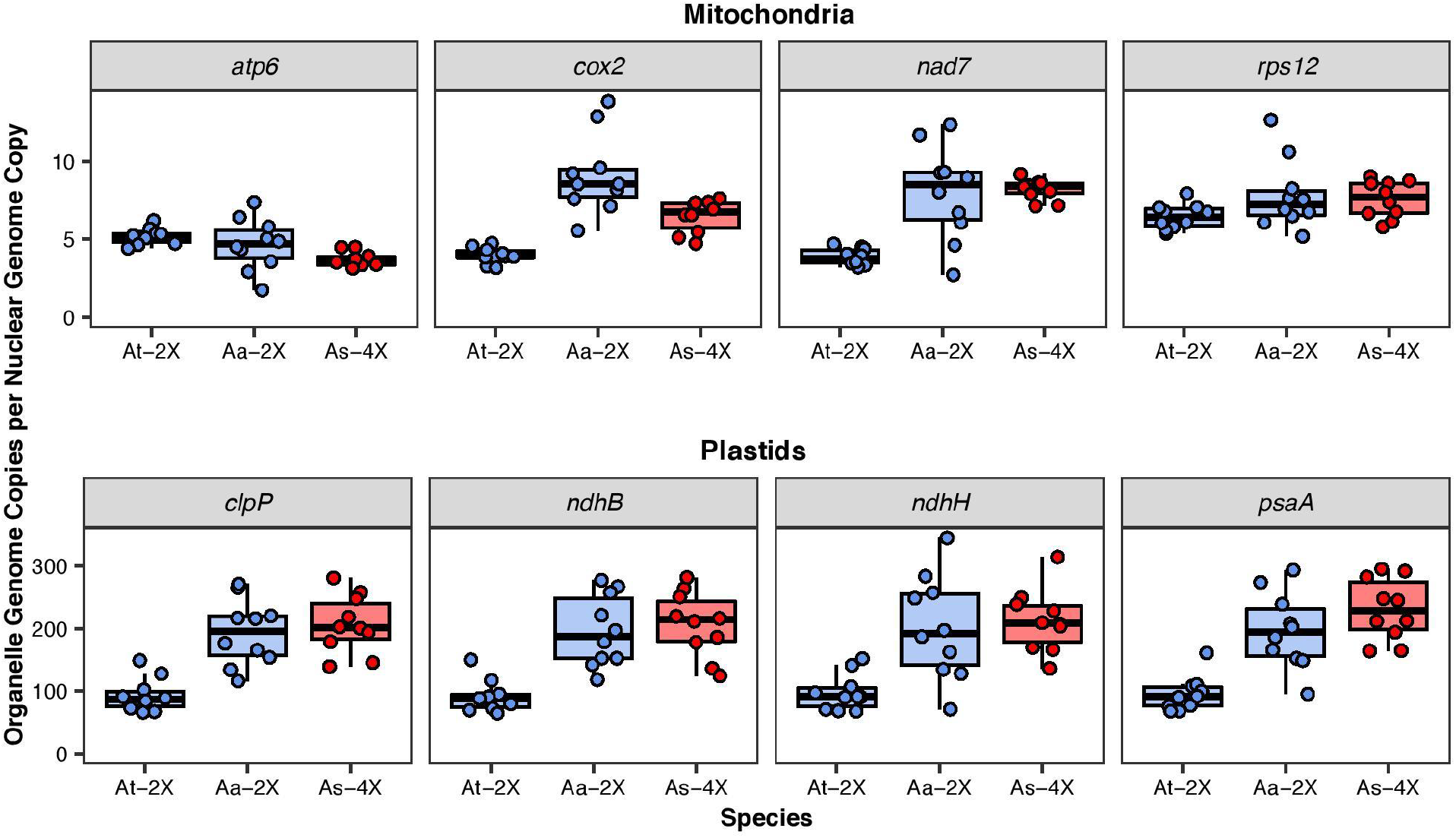
Maintenance of cytonuclear stoichiometric ratios in *Arabidopsis* allopolyploids. Organellar genome copy numbers per haploid nuclear genome copy in *Arabidopsis thaliana* (2x, blue), *Arabidopsis arenosa* (2x, blue), and *Arabidopsis suecica* (4x, red). The y-axis shows the estimated number of organellar genome copies per haploid nuclear genome copy (see Methods). Values for *atp6* and *ndhB* were divided by two to account for the fact that they are found in two-copy repeats.

**Figure S3.**
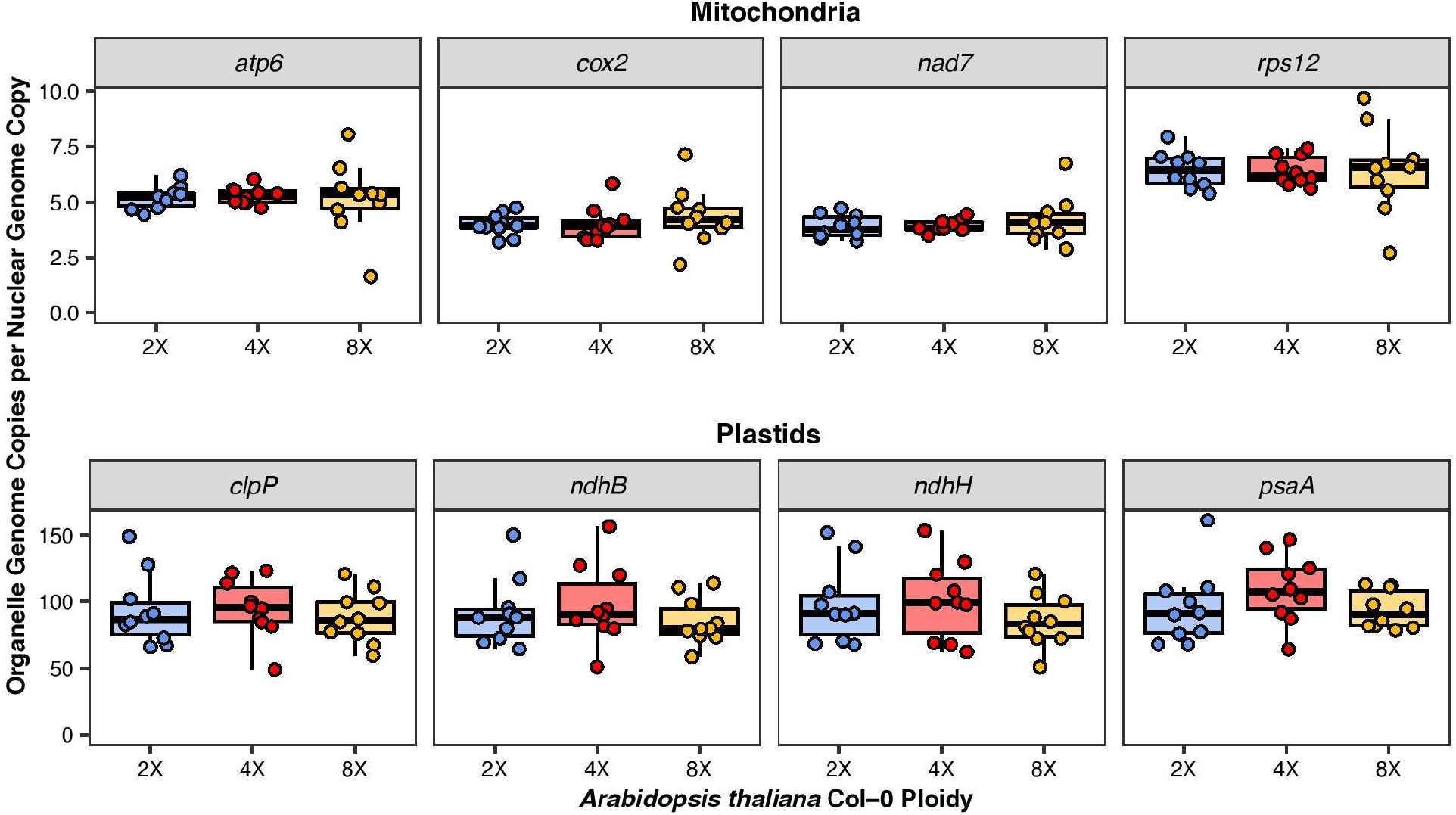
Maintenance of cytonuclear stoichiometric ratios in autopolyploid *Arabidopsis thaliana* Col-0. Organellar genome copy numbers per haploid nuclear genome copy in a synthetic polyploid series for *Arabidopsis thaliana* Col-0: 2x (blue), 4x (red), and 8x (yellow). The y-axis shows the estimated number of organellar genome copies per haploid nuclear genome copy (see Methods). Values for *atp6* and *ndhB* were divided by two to account for the fact that they are found in two-copy repeats.

**Figure S4.**
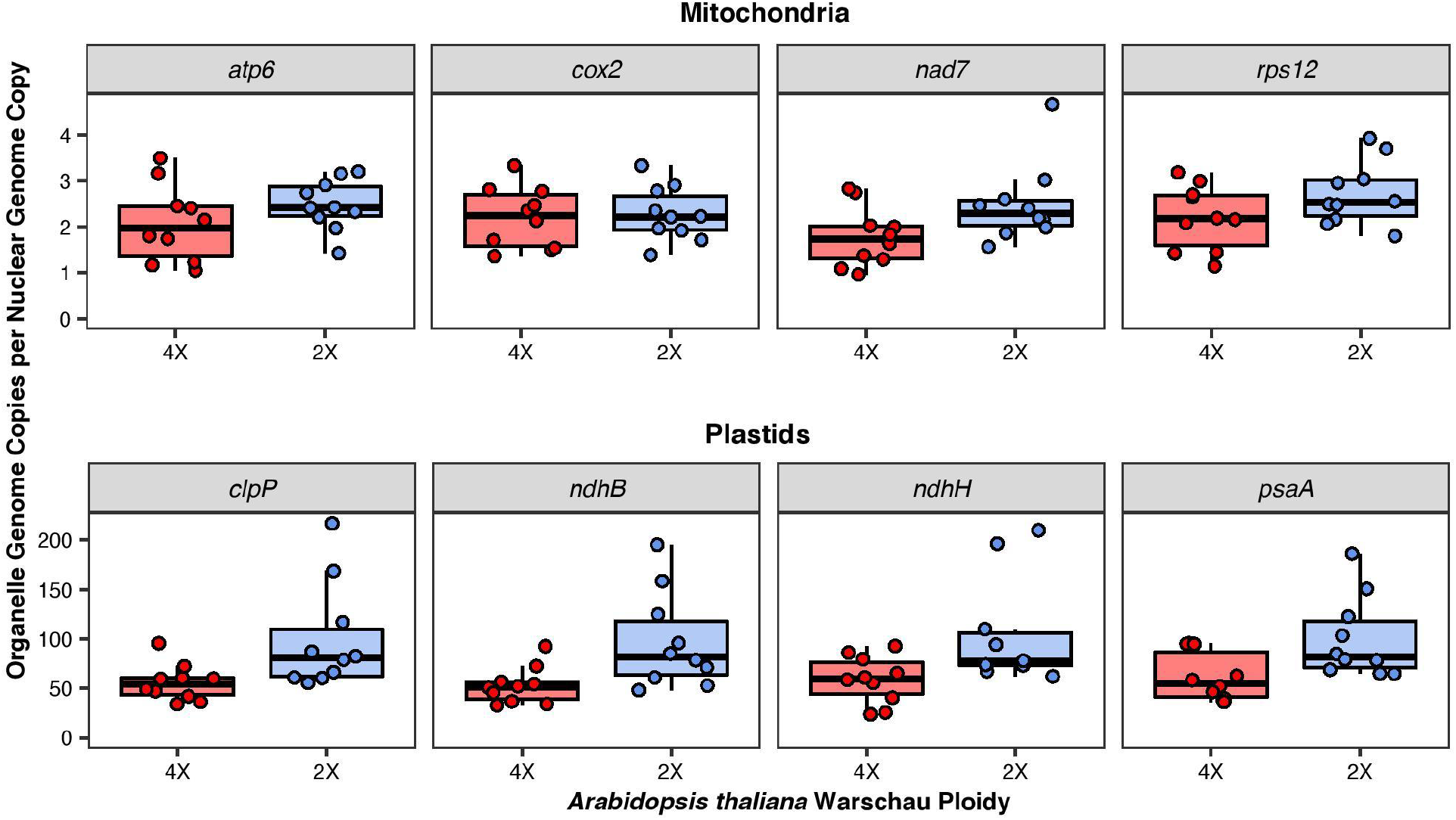
Maintenance of cytonuclear stoichiometric ratios in haploid-induced *Arabidopsis thaliana* Wa-1. Organellar genome copy numbers per haploid nuclear genome copy in *Arabidopsis thaliana* Wa-1 ecotype (4x, red) and a derived haploid induction line (2x, blue). The y-axis shows the estimated number of organellar genome copies per haploid nuclear genome copy (see Methods). Values for *atp6* and *ndhB* were divided by two to account for the fact that they are found in two-copy repeats.

## TABLES

**Table S1.** Seed sources.

| Species | Accession | Ploidy | References |
| --- | --- | --- | --- |
| *Triticum urartu* | G1799 | 2x | https://www.ars-grin.gov/ |
| *Aegilops speltoides* | As84TK097-01<br>8 | 2x | https://www.ars-grin.gov/ |
| *Triticum turgidum* | Durum No. 4 | 4x | https://www.ars-grin.gov/ |
| *Triticum aestivum* | Diehl-Mediterranean | 6x | https://www.ars-grin.gov/ |
| *Arabidopsis thaliana* | Col-0 | 2x | (Robinson *et al.*, 2018) |
| *Arabidopsis thaliana* | Col-0_4x | 4x | (Robinson *et al.*, 2018) |
| *Arabidopsis thaliana* | Col-0_8x | 8x | (Robinson *et al.*, 2018) |
| *Arabidopsis thaliana* | Wa-1_2x | 2x | (Ravi and Chan, 2010) |
| *Arabidopsis thaliana* | Wa-1 | 4x | (Ravi and Chan, 2010) |
| *Arabidopsis arenosa* | Strecno | 2x | (Lysak *et al.*, 2003)<sup>2</sup> |
| *Arabidopsis suecica* | CS 22505 | 4x | (Hanfstingl *et al.*, 1994) |

1 – National Small Grains Collection
2 – Sample was originally named *Cardaminopsis carpatica* and subsequently renamed as diploid *Arabidopsis arenosa* in 2004.

**Table S2.**
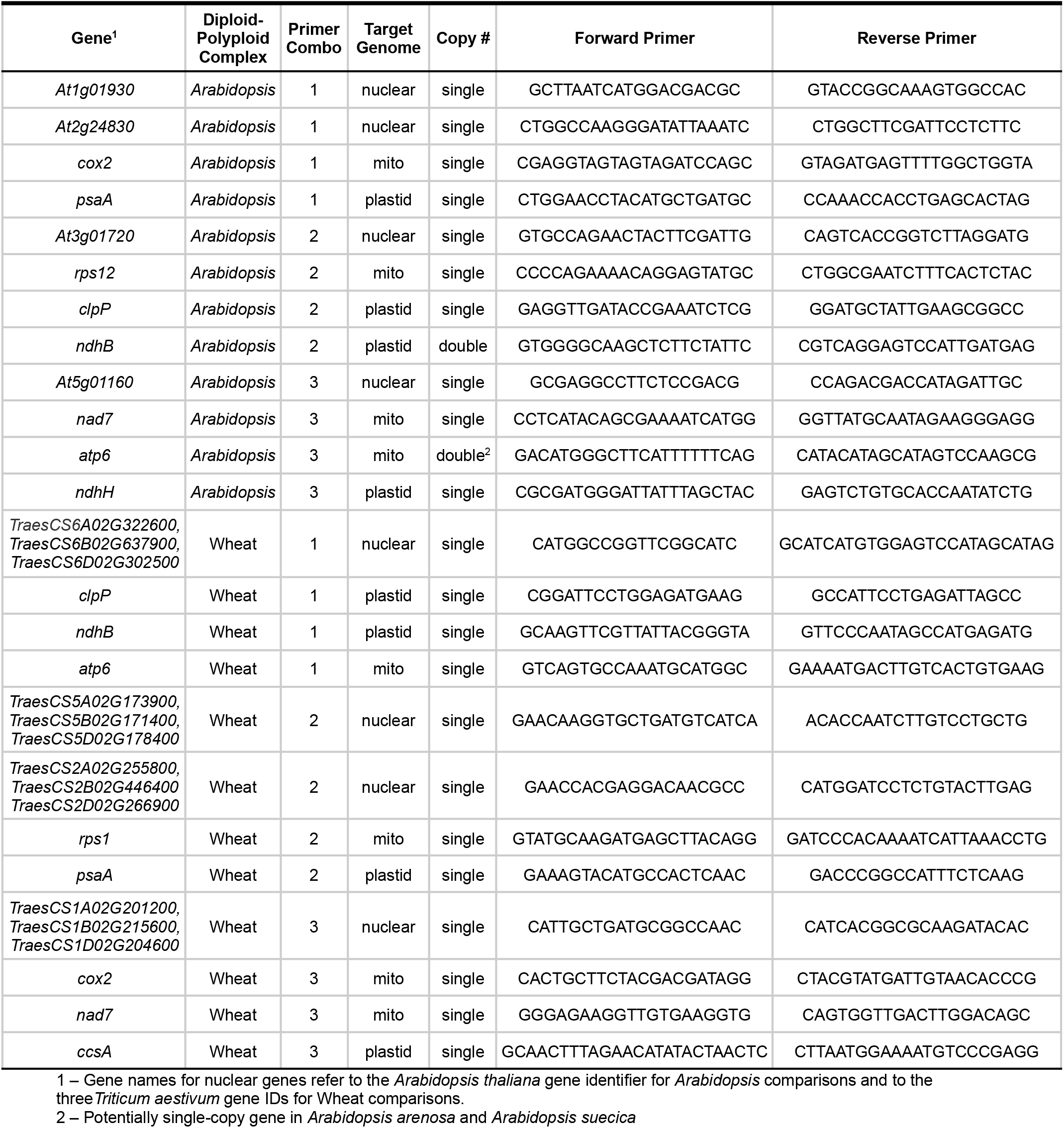
ddPCR Primers.

**Table S4.**
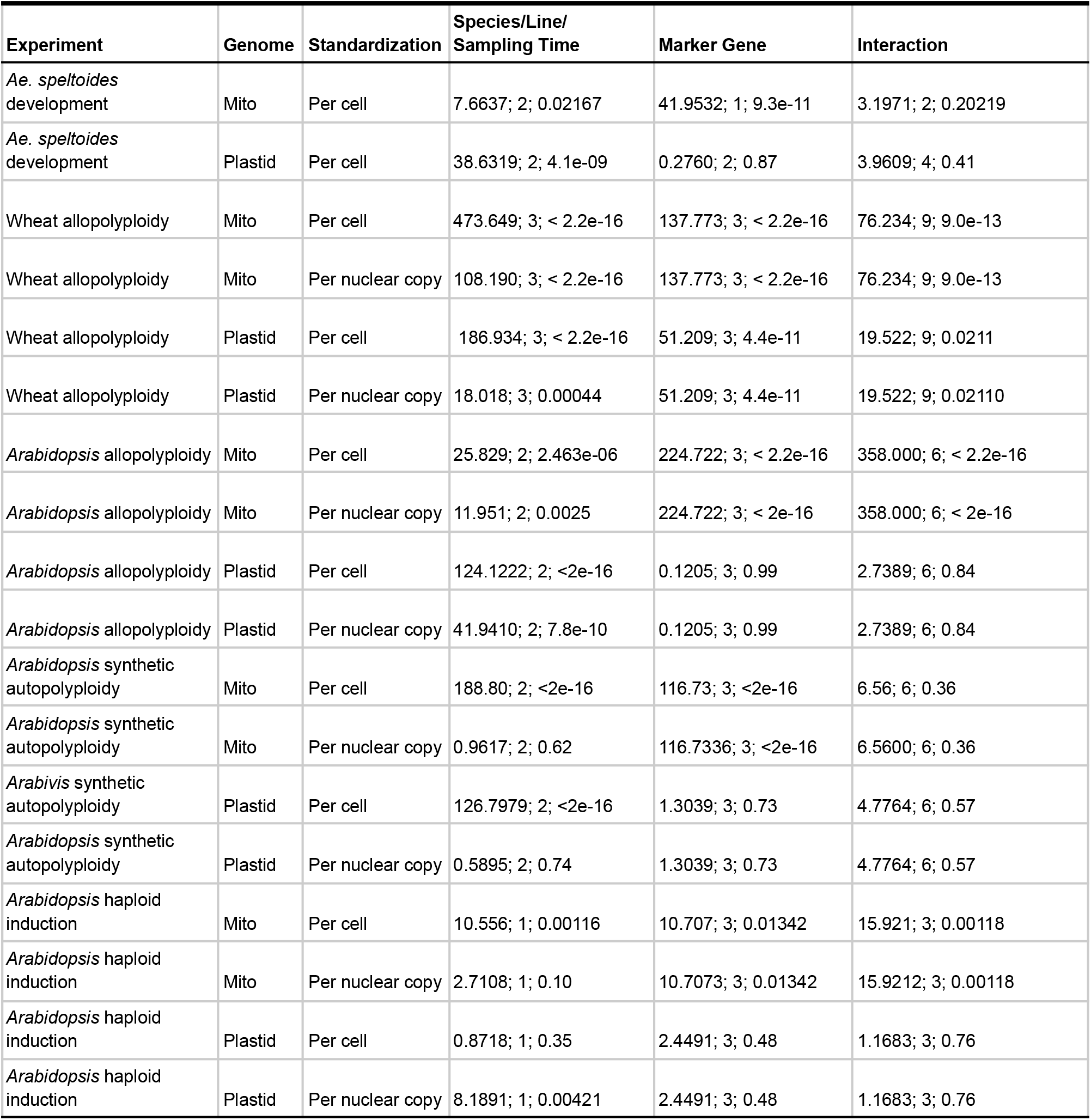
Reported data represents ***χ***^2^ values, degrees of freedom, and *P*-values for models testing for effects on organellar genome copy number.

